# The role of the diencephalon in the guidance of thalamocortical axons in mice

**DOI:** 10.1101/763201

**Authors:** Idoia Quintana-Urzainqui, P Pablo Hernández-Malmierca, James M. Clegg, Ziwen Li, Zrinko Kozić, David J Price

## Abstract

Thalamocortical axons (TCAs) cross several tissues on their journey to the cortex. Mechanisms must be in place along the route to ensure they connect with their targets in an orderly fashion. The ventral telencephalon acts as an instructive tissue, but the importance of the diencephalon in TCA mapping is unknown. We report that disruption of diencephalic development by Pax6 deletion results in a thalamocortical projection containing mapping errors. We used conditional mutagenesis to test whether these errors are due to the disruption of pioneer projections from prethalamus to thalamus and found that, while this correlates with abnormal TCA fasciculation, it does not induce topographical errors. To test whether the thalamus contains navigational cues for TCAs, we used slice culture transplants and gene expression studies. We found the thalamic environment is instructive for TCA navigation and that the molecular cues Netrin1 and Semaphorin3a are likely to be involved. Our findings indicate that the correct topographic mapping of TCAs onto the cortex requires the order to be established from the earliest stages of their growth by molecular cues in the thalamus itself.

## Introduction

A striking feature of the axonal tracts that interlink the nervous system’s component parts is the high degree of order with which they map the array of neurons in one structure onto the array of neurons in their target. Often, the order of axons at the target closely mirrors that at the source. An excellent example is the mapping of thalamic neurons onto their cerebral cortical targets via the thalamocortical pathway (Fig. 1A). Thalamic neurons located at one end of the thalamus in a dorsolateral region called the dorsal lateral geniculate nucleus (dLGN) innervate the caudal (visual) part of cortex; neurons located at the other end of the thalamus in more rostral-medial regions known as the ventrolateral (VL) and ventromedial (VM) nuclei innervate more rostral cortical regions, including motor and frontal cortex; neurons located in between - in the ventromedial posterior (VMP) nuclei - innervate central (somatosensory) cortex (Fig. 1A) (Amassian and Weiner, 1966; Bosch-Bouju et al., 2013; Jones, 2007; Tlamsa and Brumberg, 2010). The mechanisms that generate this orderly topographic mapping remain poorly understood.

**Figure 1.**
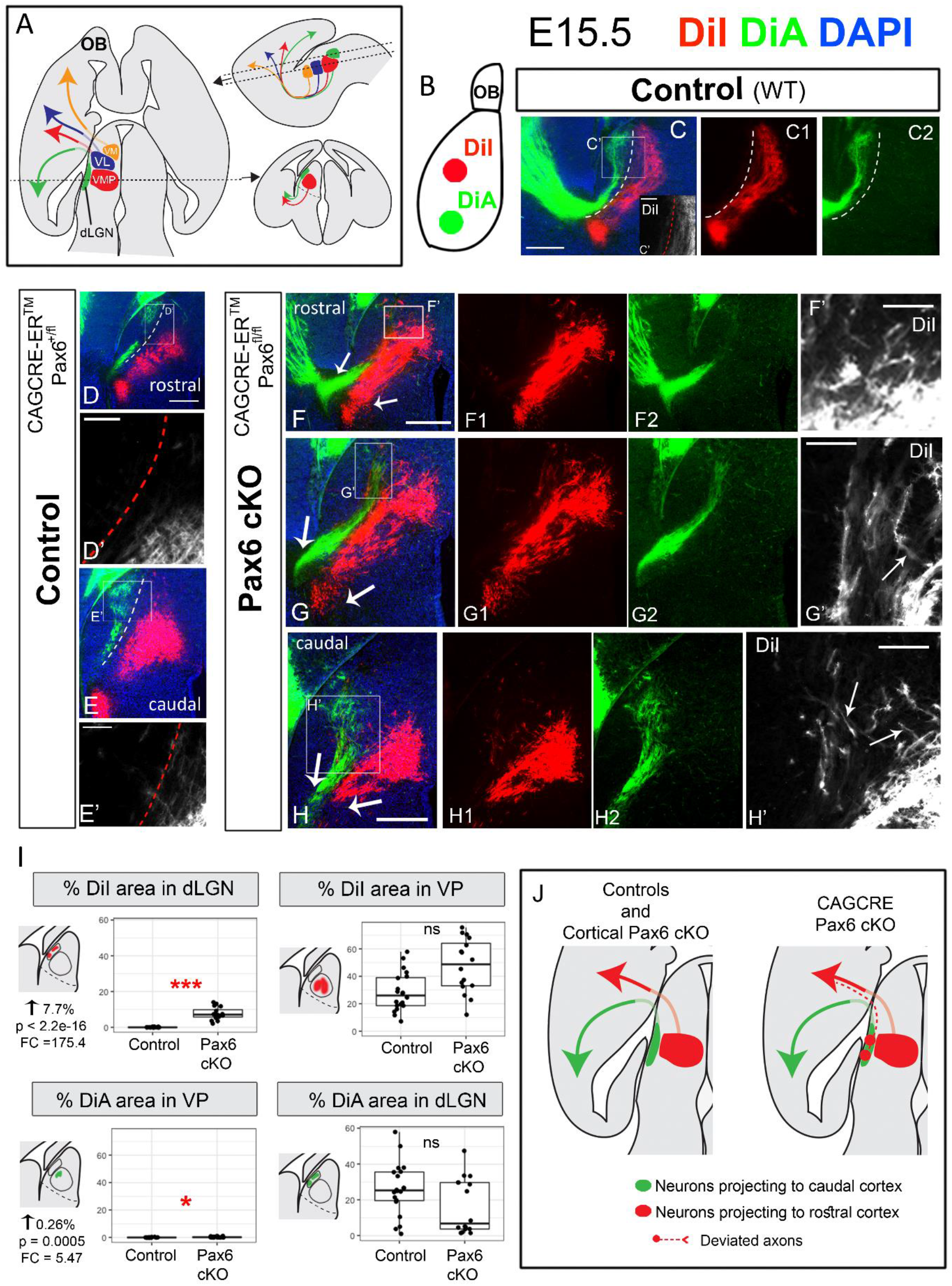
Ubiquitous conditional Pax6 deletion at E9.5 causes mapping errors in the thalamocortical connection. **A)** Schema showing how thalamic axons map to specific areas of the cortex via the thalamocortical pathway. **B)** Schematic drawing representing the cortical location where the tracers were placed at E15.5. **C-H)** Fluorescent microphotographs showing cells and axons retrogradely labelled by DiI and DiA in transverse sections. While in controls (C,E) the areas labelled in the thalamus were clearly separated (dotted lines), in CAG^CreER^ Pax6 cKOs (F-H) they visibly overlapped, indicating the existence of mapping errors in these mutants. Arrows in G,H and I indicate that the thalamocortical bundle was segregated in visual (green) and somatosensory (red) halves at the level of their exit from the diencephalon. **C’, D’, E’,F’,G’,H’)** High power images from insets in C,D,E,F,G and H, respectively, where only the DiI channel is shown. Arrows in G’ and H’ point to DiI-labeled axons showing abnormal trajectories towards medial thalamic regions. **I)** Box plots showing the area occupied by each tracer (DiI or DiA) in each defined area (dLGN or VP) in controls versus Pax6 cKOs. Seven E15.5 embryos (four controls and three Pax6 cKOs), from three different litters were analysed. For each embryo, at least five rostrocaudal sections were quantified. Each dot represents an area value of a single section. P values were calculated by fitting the data to a generalized mixed linear model. * indicates p<0.001, *** indicates p< 1.0 e-15 **J)** Schematic summary of the tract tracing results. Scale bars: 200μm (C,D,E-H); 50μm (C’,D’,E’,F’,G’,H’). dLGN= dorsal lateral geniculate nucleus, FC= Fold Change, OB= Olfactory Bulb, VL=ventral lateral nucleus, VM= ventral medial nucleus, VMP= ventral medial posterior nucleus, VP= ventral posterior, vTel= ventral telencephalon, WT= wild type.

The maintenance of order among thalamic axons as they grow is likely to contribute to the generation of orderly topographic mapping in the mature thalamocortical tract. During embryogenesis, thalamic axons exit the thalamus from about E12.5 onwards (Auladell and Hans, 2000; Braisted et al., 1999; Tuttle et al., 1999), approximately coincident with the cessation of neurogenesis in this structure (Angevine, 1970; Li et al., 2018). They then cross the adjacent prethalamus and turn laterally out of the diencephalon and into the ventral telencephalon where they traverse two consecutive instructive regions - the corridor (Lopez-Bendito et al., 2006) and the striatum - before entering the cortex. There is evidence that the maintenance of spatial order among thalamocortical axons (TCAs) crossing the ventral telencephalon requires interactions between the axons and signals released by cells they encounter in this region (Bielle et al., 2011; Bonnin et al., 2007; Braisted et al., 1999; Dufour et al., 2003; Molnár et al., 2012; Powell et al., 2008). The importance of earlier interactions within the diencephalon remains unclear.

Here, we tested the effects of mutating the gene for the Pax6 transcription factor, which is essential for normal diencephalic patterning (Caballero et al., 2014; Clegg et al., 2015; Parish et al., 2016; Pratt et al., 2000; Stoykova et al., 1996; Warren and Price, 1997), on the topographic mapping of TCAs onto the cortex. Pax6 starts to be expressed in the anterior neural plate well before TCAs start to form (Walther and Gruss, 1991). As the forebrain develops from the anterior neural plate, Pax6 expression becomes localized in (i) cortical progenitors that generate the target neurons for TCAs, (ii) diencephalic (thalamic and prethalamic) progenitors and (iii) prethalamic (but not thalamic) postmitotic neurons (Quintana-Urzainqui et al., 2018; Stoykova et al., 1996; Warren and Price, 1997). We discovered that deletion of Pax6 from mouse embryos at the time when thalamic axons are starting to grow results in the development of a thalamocortical projection containing mapping errors. Axons from dorsolateral thalamus are misrouted medially and end up projecting abnormally rostrally in the cortex. We went on to explore the reasons for this defect.

We first used conditional mutagenesis to test whether misrouting is due to the loss of Pax6 from prethalamic neurons, since previous work has shown that (i) Pax6 is not required in the cortex for normal TCA topography (Piñon et al., 2008) and (ii) Pax6 is neither expressed nor required autonomously by thalamic neurons for them to acquire the ability to extend axons to the cortex (Clegg et al., 2015). We found that while loss of Pax6 from a specific set of prethalamic neurons prevented them developing their normal axonal projections to thalamus and resulted in the abnormal fasciculation of thalamic axons, it did not cause TCAs to misroute. This suggested that the thalamus itself contains important navigational cues for TCAs. We used slice culture transplants and gene expression studies to show (i) that the thalamic environment is indeed instructive for TCA navigation and (ii) to identify molecular changes within the thalamus that likely cause the disruption in TCA topography observed upon Pax6 deletion. Our findings indicate that the normal topographic mapping of TCAs requires that order be established and maintained from the earliest stages of their growth by molecular cues in the thalamus itself.

## Results

### Thalamocortical topography is disrupted in *CAG^CreER^* but not in *Emx^CreER^* Pax6 conditional knockouts

Previous studies have shown that constitutive loss of Pax6 function causes a total failure of TCA development, which is hypothesized to be a secondary consequence of anatomical disruption at the interface between the diencephalon and the telencephalon (Clegg et al., 2015; Georgala et al., 2011; Jones et al., 2002). No such failure occurs if Pax6 is deleted conditionally after this anatomical link is formed (Clegg et al., 2015). We first assessed whether delayed ubiquitous Pax6 deletion, induced in *CAG^CreER-TM^ Pax6^fl/fl^* embryos (referred to here as *CAG^CreER^* Pax6 cKOs), disrupts the topography of TCA connections. We induced Cre recombinase activation by tamoxifen administration at E9.5 which caused Pax6 protein loss in *CAG^CreER^* Pax6 cKOs from E11.5 onwards (Quintana-Urzainqui et al., 2018), which is when the generation of most thalamic neurons is starting (Li et al., 2018) and before many TCAs have begun to grow (Auladell and Hans, 2000; López-Bendito and Molnár, 2003). Diencephalic progenitor domains in *CAG^CreER^* Pax6 cKOs are fully recognizable (Quintana-Urzainqui et al., 2018) and patterning seems largely unaffected in the thalamus (Fig. S1). We used both wild type and *CAG^CreER^ Pax6^fl/+^* littermate embryos as controls since the latter express normal levels of Pax6 protein (see Methods; Caballero et al., 2014; Manuel et al., 2015).

We inserted two different axonal tracers in two cortical areas in E15.5 fixed brains. DiA was placed in the visual (caudal) cortex while DiI was placed in the somatosensory (more rostral) cortex (Fig. 1B). In controls (both wild type and *Pax6^fl/+^*), DiA retrogradely labelled cells in dorsolateral thalamic areas (dLGN; green labelling in Fig. 1C-E), while DiI labelled cells in ventromedially-located thalamic regions, identified as ventral-posterior thalamic nucleus (VP) (red labelling in Fig. 1C-E). Labelling of these two thalamic regions was clearly separated in all cases (indicated by dotted line in Fig. 1C-E). In *CAG^CreER^ Pax6* cKOs, however, the two labelled populations overlapped (Fig. 1F-H). In these mutants, the distribution of the DiA-labelled thalamic cells (from caudal cortical injections) was not obviously changed with respect to controls. However, the DiI-labelled thalamic cells (projecting to more rostral cortical areas) showed a much wider distribution than in controls and expanded to lateral thalamic areas (compare Fig.1 C-E,C’-E’ versus F-H,F’-H’), even overlapping with DiA stained cells at the dLGN (Fig. 1 G,H,G’,H’). To quantitate this, we measured the area occupied by DiI and DiA within each nucleus of interest in controls versus *CAG^CreER^ Pax6* cKOs (in transverse E15.5 sections from three different litters: four controls, three cKOs, five sections per embryo). We defined the dLGN and VP nucleus as regions of interest (ROI) blind to DiI/DiA labelling and measured the percentage of each ROI occupied by DiI and DiA (for details see Methods). We found highly significant increase (from virtualy 0% to 7.7%) of the area occupied by DiI in the dLGN in mutants compared to controls (Fig. 1I). We also found a significant increase in DiA in VP nucleus, although its magnitude was very small (0.26%) (Fig. 1I). Areas occupied by DiI in VP or DiA in dLGN were not significantly altered. This analysis shows that *CAG^CreER^ Pax6* cKOs display topographic errors, the main one being the misrouting of axons from dLGN neurons towards abnormally rostral cortical areas in the *CAG^CreER^ Pax6* cKOs.

Since Pax6 is expressed both in the cortex and diencephalon during TCA development, the mapping defects described above might have been due to the loss of Pax6 from the cortex. This was unlikely because a previous study showed that Pax6 is not required in the cortex for the establishment of proper topographical thalamocortical connections (Piñon et al., 2008). To confirm this, we used a cortex-specific, tamoxifen-inducible Cre line (*Emx1^CreER^*). We administered tamoxifen at E9.5, which results in a near-complete loss of cortical Pax6 between E11.5-12.5 (Georgala et al., 2011; Mi et al., 2013), and performed DiI/DiA labelling at E15.5, following the same experimental design described above for the *CAG^Cre^* line. We found that the two retrogradely-labelled populations did not overlap in controls (*Emx1^CreER^ Pax6^fl/+^*) or in mutants (*Emx1^CreER^ Pax6^fl/fl^*) (Fig. S2A,B), suggesting that the defects of TCA mapping found in the *CAG^CreER^ Pax6* cKOs were not attributable to cortical abnormalities.

To define the anatomical region where thalamic axons probably deviated from ordered growth, we examined the TCA bundle in *CAG^CreER^ Pax6* cKOs. This bundle was ordered and segregated into rostral/somatosensory (DiI) and caudal/visual (DiA) halves at its point of exit from the prethalamus and entry into the ventral telencephalon (arrows in Fig. 1 F-H), which indicated that the misrouting of lateral TCAs from dorsolateral thalamus might happen before this point, i.e. within the diencephalon.

### TCAs fasciculate prematurely as they cross the prethalamus in *Pax6* conditional mutants

Within the diencephalon, the first structure that thalamic axons encounter as they leave the thalamus is the prethalamus, and its neurons normally express high levels of Pax6. Therefore, we investigated whether the defects of TCA mapping in *CAG^CreER^ Pax6* cKOs might arise from a disordered growth of thalamic axons through the prethalamus. As a first step, we examined the effects of Pax6 deletion on the behaviour of thalamic axons as they cross the prethalamus.

Since the neural cell adhesion molecule L1CAM (L1) is expressed in TCAs (Fukuda et al., 1997; Ohyama et al., 2004), we examined the distribution of L1-positive thalamic axons at E13.5 in transverse and sagittal sections through the prethalamus. In controls, axons emerging from the thalamus converge progressively as they cross the prethalamus (Fig. 2A-E) to subsequently form a single thalamocortical bundle that turns laterally and exits the prethalamus (arrows in Fig. 2A,B,E). We found that in *CAG^CreER^ Pax6* cKOs (Fig. 2F-L), thalamic axons prematurely converge into larger bundles as soon as they cross the thalamic-prethalamic boundary (empty arrows in Fig. 2G-I,L).

**Figure 2.**
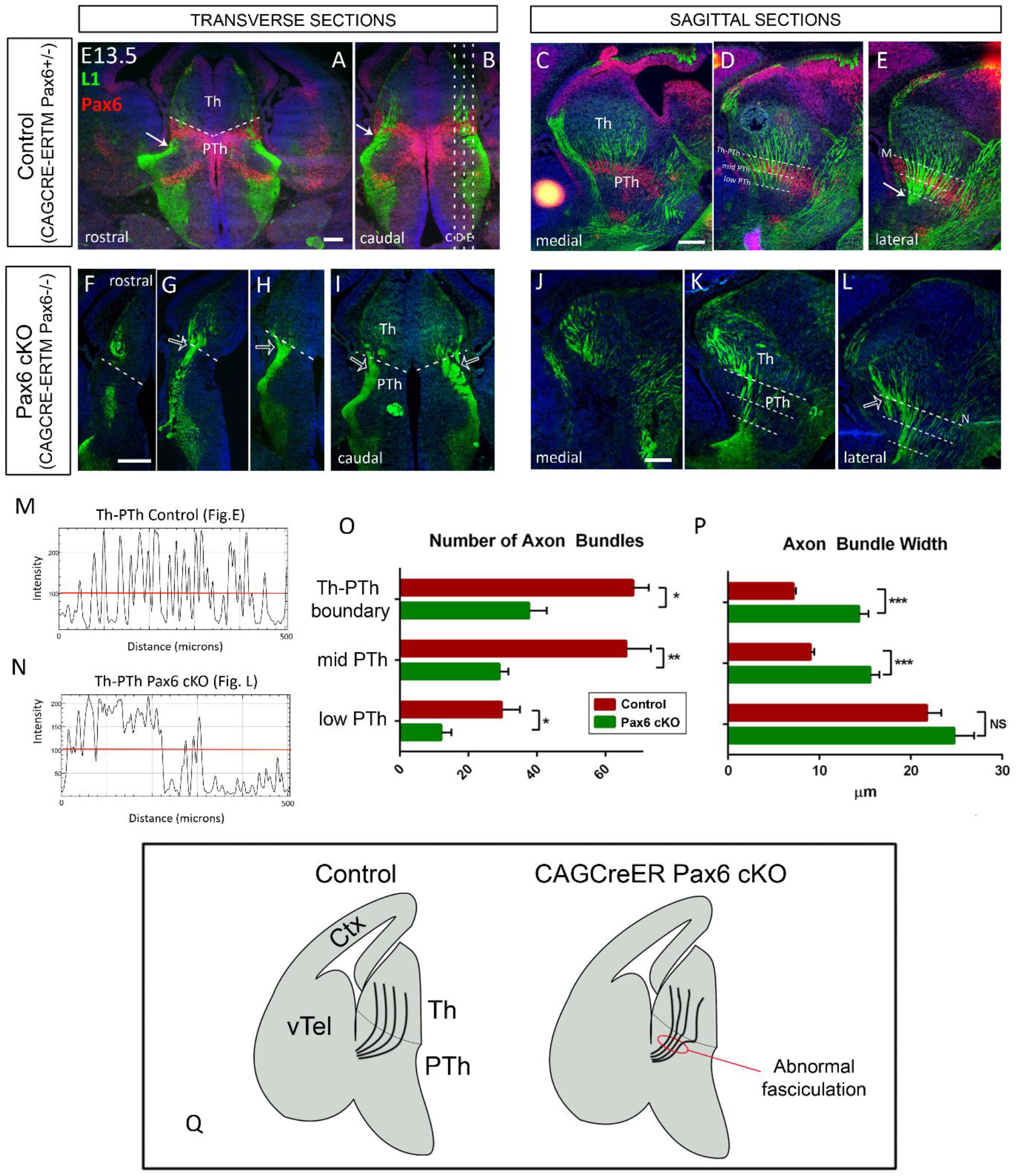
Thalamic axons exhibit abnormal fasciculation as they cross the prethalamus in CAG^CreER^ Pax6 cKOs. **A-L)** Transverse (A,B, F-I) and sagittal (C-D, J-L) sections showing immunofluorescence for L1 and Pax6 in controls (A-E) and CAG^CreER^ Pax6 cKOs (F-L) at E13.5. White arrows in A,B and E show the point at which the thalamocortical tract forms a single bundle and exits the diencephalon in controls. Empty arrows in G-I and L point to thalamic axons converging prematurely into big bundles as soon as they cross the thalamic-prethalamic boundary in Pax6 cKOs. Discontinuous lines in A, F-I mark the thalamic-prethalamic boundary. Discontinuous lines in B mark the level of sections in C, D and E. Discontinuous lines in D,E,K and L show the position of the lines used to quantify the number and width of bundles crossing the prethalamus. **M,N)** Examples of the measurements taken for the quantifications of the number and width of axon bundles crossing each line. Individual bundles were identified as single L1-positive structure above a consistent intensity threshold (red lines). **O,P)** Graphs showing the quantification of the number (O) and width (P) of axon bundles crossing each reference line and the statistical significance. Representation of means +/- SEM, n=3 (three embryos analysed from three different litters, five slices per embryo), where *, ** and *** stand for a p-value ≤ 0.05, 0.01 and 0.001 respectively after two-tailed unpaired Student’s t-test and N=3. O) We found a significant decrease in the number of bundles crossing all three checkpoints (Th-PTh boundary: p value=0.046, t=2.86, df=4; Mid Pth: p value=0.0085, t=4.82, df=4; low-PTh: p-value=0.012, t=4.39, df=4). P) We detected a big increase in axon bundle width at the Th-PTh border (p-value<<0.001, t=9.12, df=443) and the mid-PTh (p-value<< 0.001, t=7.21, df=350), but no significant change at the low-PTh. **Q)** Schematic summary of the results showing how thalamic axons undergo abnormal and premature fasciculation when crossing the prethalamus in CAG^CreER^ Pax6 cKOs. Scale bars: 200μm. Ctx= cortex, Pth= prethalamus, Th= thalamus, vTel= ventral telencephalon.

To obtain a quantitative measurement of this observation we positioned three equally-spaced lines across different diencephalic levels in sagittal sections (see Methods; lines represented in Fig. 2D,E,K,L). We used Fiji software (Schindelin et al., 2012) to quantify the number of axon bundles crossing each line and the width of each individual bundle. We recognized individual bundles as each single L1-positive structure above a consistent intensity threshold (red lines in Fig. 2M,N). We found a significant decrease in the number of bundles crossing all three checkpoints (Fig. 2O). Axon bundle width strikingly increased at the Th-PTh border and the mid-PTh, with no significant change at the low-PTh checkpoint line (Fig. 2P) (see figure legend for statistical details). These data indicate that, in the absence of Pax6, TCAs begin to fasciculate prematurely in their route, forming bigger and fewer bundles as they cross the prethalamus (Fig. 2Q). We next tested the role of the prethalamus in TCA formation and the potential establishment of their topography.

### Loss of prethalamic pioneer axons correlates with abnormal TCA fasciculation but not with changes in topography

The prethalamus has been proposed to host a population of neurons that extend axons to the thalamus which act as “pioneer guides” for TCA navigation (Price et al., 2012; Tuttle et al., 1999). We assessed whether this population is disrupted by Pax6 loss from the prethalamus since, if it is, this might provide an explanation for phenotypes described above.

From E9.5 on, most cells in the prethalamus express, or are derived from cells that expressed, *Gsx2*. We used a *Gsx2^Cre^* line (Kessaris et al., 2006) carrying an EGFP Cre reporter (Sousa et al., 2009) to visualize neurons and axons belonging to the *Gsx2* lineage and we observed that prethalamic Pax6-expressing cells are included within the location of the *Gsx2* lineage prethalamic population (Fig. 3A,B). *Zic4* is also expressed by some prethalamic cells, with an onset of expression similar to that of *Gsx2* (about E9.5; Gaston-massuet et al., 2005), and most diencephalic *Zic4* lineage cells express and require Pax6 for their normal development (Li et al., 2018). Using a *Zic4^Cre^* line (Rubin et al., 2011) we observed that prethalamic neurons derived from *Zic4* lineage were located in a narrow band close to the thalamic-prethalamic border (Fig. 3C-D). We assessed whether these prethalamic populations normally send axons to the thalamus.

**Figure 3.**
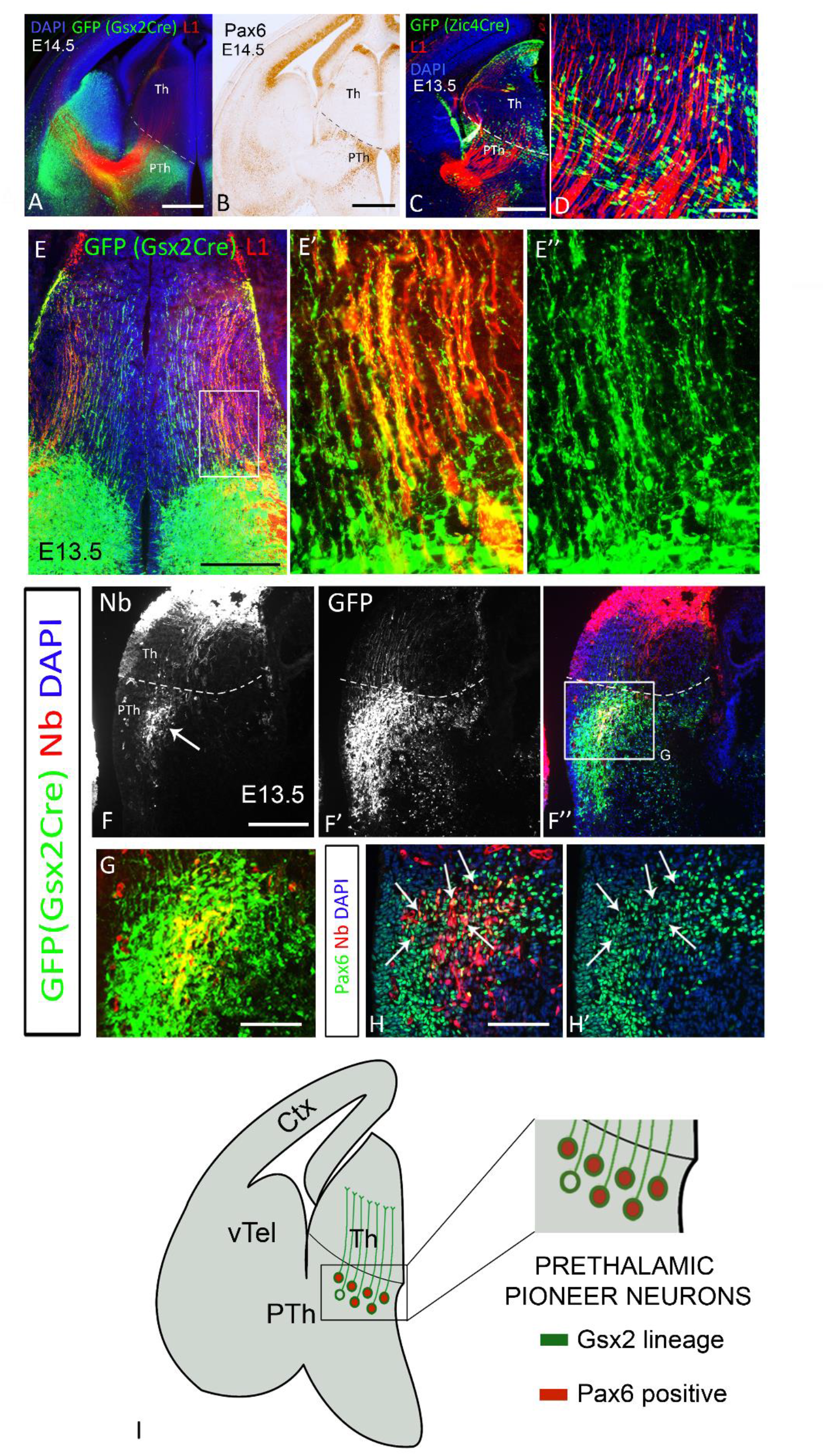
Prethalamic pioneer axons belong to the Gsx2 lineage and express Pax6. **(A)** EGFP reporter of Cre recombinase activity and L1 staining in Gsx2^Cre^ embryos at E14.5. **B)** Immunohistochemistry showing Pax6 expression **C)** EGFP reporter of Cre activity in Zic4^Cre^ embryos at E13.5 and thalamocortical axons expressing L1. **D)** High power image showing the absence of axonal projections from Zic4-lineage prethalamic cells towards the thalamus. **E-E’’)** Neurons belonging to the Gsx2 lineage extend parallel projections across the thalamus in close apposition to L1-positive thalamocortical axons. E’ and E’’ are a higher magnification of the area framed in E. **F-H)** Injection of the tracer Neurobiotin in the thalamus at E13.5 resulted in the retrograde labelling of neurons in the prethalamus (arrow in F). F-G: Double labelling of GFP and Neurobiotin shows that prethalamic labelled neurons belong to the Gsx2. H,H’: Parallel section of F” processed for Pax6 immunoshistochemistry combined with Neurobiotin visualization showing that most prethalamic neurobiotin-positive neurons are also Pax6 positive (arrows). **I)** Schematic summary. Scale bars: 500μm (A,B,C,E), 250μm (F), 100μm (D, E’,G,H). Ctx= cortex, Nb= Neurobiotin, Pth= prethalamus, Th= thalamus, vTel= ventral telencephalon. All descriptions where observed in at least three embryos from three different litters.

*Gsx2*-lineage GFP-positive axons extended throughout the thalamus forming ordered and parallel projections (Fig. 3E) and running in close apposition to L1-positive TCAs (Fig. 3E’,E’’) from E12.5 onwards (Fig. S3). By contrast, *Zic4*-lineage prethalamic cells did not project axons to the thalamus (Fig. 3D), indicating that prethalamic pioneer axons arise from *Gsx2*-lineage and not from *Zic4*-lineage prethalamic cells.

Since *Gsx2* is also expressed in the ventral telencephalon (Fig. 3A), and ventral telencephalic neurons are known to project to the thalamus (López-Bendito and Molnár, 2003; Métin and Godement, 1996; Molnár et al., 1998), there was a possibility that *Gsx2^Cre^* lineage axons innervating the thalamus actually originated from ventral telencephalic neurons. To confirm the existence of *Gsx2*-lineage prethalamic neurons projecting to the thalamus we injected the neuronal tracer Neurobiotin^™^ in the thalamus of E13.5 *Gsx2^Cre^* embryos and successfully labelled prethalamic neurons (arrow in Fig. 3F). Neurobiotin^™^-positive cells were GFP-expressing *Gsx2* lineage (Fig. 3F-F’’,G) and most of them also expressed Pax6 (arrows in Fig. 3H,H’; see summary in Fig. 3I). (Note that individual injections each involved only subregions of the thalamus, explaining why each one only labelled a discrete subset of the prethalamic neurons projecting to the thalamus). This experiment confirmed that *Gsx2*-lineage cells in the prethalamus project to the thalamus.

Having established that pioneer prethalamic axons belong to the *Gsx2* lineage and express Pax6 we next aimed at disrupting their formation by conditionally deleting Pax6 in *Gsx2*-lineage cells. We crossed mice carrying the floxed *Pax6* allele and EGFP Cre reporter with the *Gsx2^Cre^* line. Pax6 conditional deletion in *Gsx2*-lineage cells (*Gsx2^Cre^ Pax6* cKOs) caused a visible reduction in the number of GFP-positive axons projecting from prethalamus to thalamus in E12.5, E13.5 and E14.5 embryos (Fig. 4A-F). To confirm that the prethalamic axons that were lost in *Gsx2^Cre^*; *Pax6^loxP/loxP^* embryos were Pax6-expressing, we used the DTy54 YAC reporter allele to express tauGFP in cells in which the *Pax6* gene is active, irrespective of whether it is mutant or not (Tyas et al., 2006). Whereas there were many tauGFP-labelled axons running from prethalamus to thalamus in controls, there were very few in experimental embryos (Fig. 4G-J).

**Figure 4.**
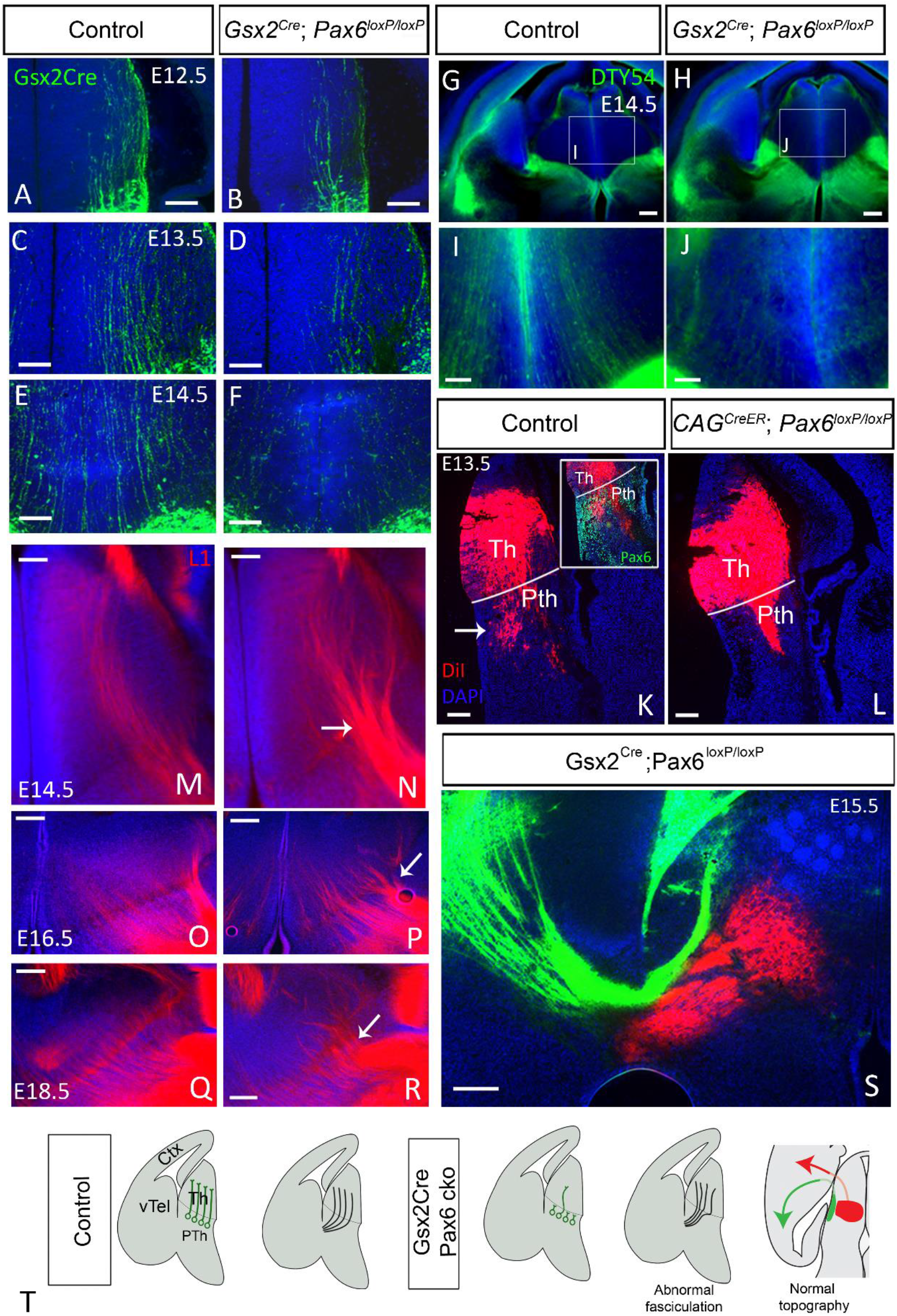
Disruption of prethalamic pioneer axons in Gsx2^Cre^ Pax6 cKOs causes abnormal fasciculation but not changes in topography. **A-F)** Prethalamic pioneer axons visualized by EGFP reporter in Gsx2^Cre^ embryos are reduced in Gsx2^Cre^ Pax6 cKOs at E12.5 (A,B), E13.5 (C,D) and E14.5 (E,F) **G-J)** Use of the DTy54 YAC reporter allele, which labels cells in which the *Pax6* gene is active, also reveals a reduction in prethalamic pioneer axons crossing the thalamus. **K,L)** Injection of the tracer DiI in the thalamus of E13.5 retrogradelly labels cell bodies in the prethalamus of controls (arrow in K) but not in CAG^CreER^ Pax6 cKOS (L). Inset in K is a fluorescent immunohistochemistry for Pax6 combined with DiI visualization showing that the DiI-labelled prethalamic neurons are expressed in the Pax6 positive area. **M-R)** Immunofluorescence for L1 shows formation of abnormal big bundles of thalamocortical axons crossing the prethalamus in Gsx2^Cre^ Pax6 cKOs (arrow in N,P,R) with respect to controls (M,O,Q) at E14.5 (M,N), E16.5 (O,P) and E18.5 (Q,R). **S)** DiI and DiA injection in the cortex of Gsx2^Cre^ Pax6 cKO embryos shows that the topography of TCAs is not affected in these mutants. **T)** Schematic summary. Scale bars: 250μm (G,H,S), 100 μm (A-F, I-P). Ctx= cortex, Pth= prethalamus, Th= thalamus, vTel= ventral telencephalon. All phenotypes where observed in at least three embryos from three different litters, unless otherwise stated.

DiI placed in the thalamus of E13.5 *CAG^CreER^* controls (*CAG^CreER^ Pax6^fl/+^*) retrogradely labelled a prethalamic population (arrow in Fig. 4K). In the absence of *Pax6* (*CAG^CreER^ Pax6* cKOs), no prethalamic cell bodies were labelled by thalamic DiI injection (Fig. 4L), providing further evidence that the prethalamic pioneer population does not form correctly when Pax6 is deleted. Overall, our results show that prethalamic pioneer axons originating from *Gsx2*-lineage cells both express and require Pax6 to develop normal connections with the thalamus (Fig. 4T).

We then studied the TCAs of *Gsx2^Cre^ Pax6* cKOs. Similar to the phenotype described in *CAG^CreER^ Pax6* cKOs, Pax6 deletion in *Gsx2* lineage caused abnormal premature fasciculation of axons crossing the thalamic-prethalamic border, as evidenced by L1 immunohistochemistry (Fig. 4M-R). However, unlike *CAG^CreER^ Pax6* cKOs, *Gsx2^Cre^ Pax6* cKOs did not show abnormal topographical projections, with no obvious overlap between thalamic retrogradely-labelled populations after cortical DiA and DiI placement in caudal and more rostral cortex respectively (Fig. 1B; Fig. 4S). We conclude that, while prethalamic pioneer axons may play a role in avoiding premature TCA fasciculation, they are not required for the establishment of accurate thalamocortical topographic mapping (Fig. 4T).

### Evidence for the importance of navigational cues in the thalamus itself

We next considered the potential importance of thalamic factors in the establishment of thalamocortical topographic order. Our results above indicate that thalamic axons might have deviated from their normal trajectories before they exited the thalamus in *CAG^CreER^ Pax6* cKOs (arrows mark deviant axons in Fig. 1 G’,H’; no such axons were observed in the controls, Fig. 1C’,D’,E’). A misrouting of TCAs in the thalamus was also evident with L1 staining (Fig. 5 A,B). The largest collections of deviant axons were observed projecting from lateral to medial thalamic regions (arrow in Fig. 5B), suggesting that the loss of Pax6 had disrupted normal navigational mechanisms operating within the thalamus.

**Figure 5.**
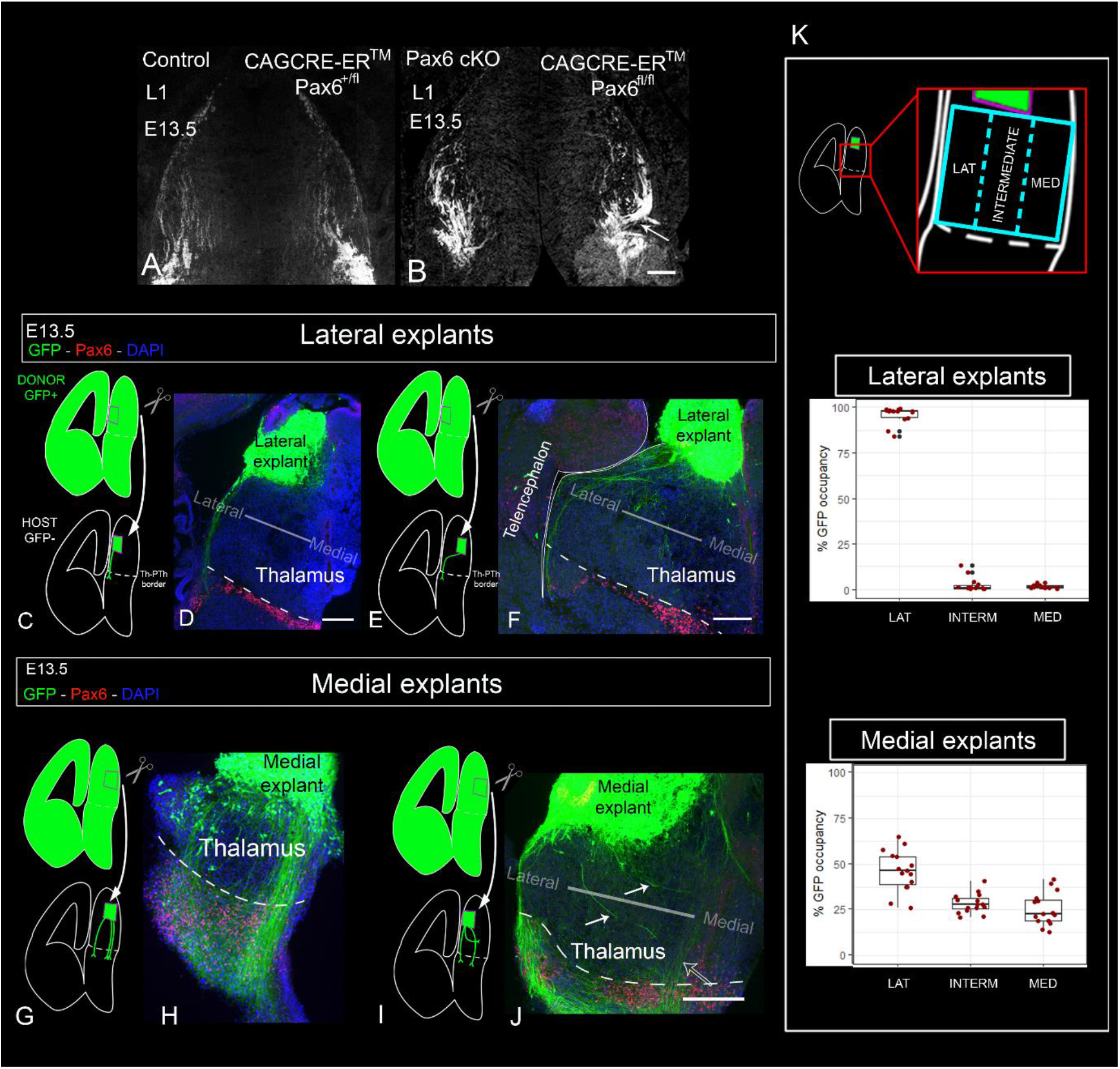
Subsets of thalamocortical axons show a preference to navigate through different parts of the thalamus. **A,B)** Immunohistochemistry for L1 shows that thalamocortical axons exhibit disorganized and erroneous trajectories in the thalamus of E13.5 CAG^CreER^ Pax6 cKOs. Arrow in B points to one big bundle deviating from lateral to medial thalamus. **C-J)** Schemas and images of the transplants using lateral (C-F) or medial (G-J) donor thalamus and grafted into lateral (C,D,G,H) or medial (E,F,I,J) host tissue. Images show immunofluorescence for GFP and Pax6. White arrows in J show medial axons turning from lateral to medial regions of the thalamus.. **K)** Quantification of the navigation of GFP-positive axons through three sectors of the thalamus shows that lateral axons have a strong preference to navigate through lateral thalamus while medial axons show a more variable response. We quantified six thalamic graft experiments with embryos from three different litters (three using lateral explants and three using medial explants). Box plots show the percentage of GFP-positive axons detected on each sector of the thalamus. Five rostrocaudal sections were quantified per explant. Each dot represents an area value of a single section. Scale bars: 100μm. Ctx= cortex, Pth= prethalamus, Th= thalamus, vTel= ventral telencephalon.

We looked for evidence that thalamic axons are actively guided through the normal thalamus by using *in vitro* slice culture transplants to assess the effects of repositioning lateral or medial thalamic neurons on the routes taken by their axons. We grafted thalamic slice explants from E13.5 GFP-positive donor embryos into GFP-negative host slices and cultured them for 72 hours to allow thalamic axons to regrow and navigate through the host environment. The donor grafts were positioned so that their axons had to traverse at least 200μm of host thalamic tissue before encountering the Th-PTh boundary, allowing us to assess how the host thalamic tissue affected the trajectory of the axons emerging from the donor tissue. We isolated donor explants from either lateral or medial thalamus, and grafted them either medially or laterally into host thalamus (Fig. 5C,E,G,I).

We found that axons from lateral thalamic explants showed a strong preference to follow a lateral trajectory, while axons from medial explants extended much broadly throughout the thalamus. In both cases, this effect was irrespective of whether they were grafted laterally or medially (Fig. 5C-J). To quantitate this, we divided the thalamus in three equal sectors (lateral, intermediate, medial; Fig 5K) and measured the area occupied by GFP-positive elements in each of the sectors. We quantified six explants (three lateral and three medial, 5 sections per explant) from three different litters. In average, 95.7% of axons from lateral explants navigated through lateral-most area (Fig 5.K), while medial explants extended axons more evenly across the three regions (45.8% lateral, 31.8% intermediate and 22.4% medial; Fig. 5K). Moreover, we observed that when lateral axons were confronted with medial host thalamus, most made a sharp turn towards lateral positions before heading towards the prethalamus (Fig. 5E,F). When medial explants were grafted laterally many of their axons turned medially (arrows in Fig. 5J).

These results indicated that different subsets of thalamic axons exhibit different chemotactic responses to the thalamic environment and therefore that thalamic axons are actively guided by mechanisms operating within the thalamus itself. To gain further insight into what these mechanisms might be, we went on to examine the expression of guidance molecules in the normal thalamus and in thalamus from which Pax6 has been removed.

### Axon guidance molecule expression in normal and Pax6 deficient thalamus

Semaphorin 3a (Sema3a) and Netrin 1 (Ntn1) are secreted guidance molecules whose complementary gradients of expression in the ventral telencephalon are key for the correct establishment of topographical connections between thalamus and cortex (Bielle et al., 2011; Braisted et al., 1999; Molnár et al., 2012; Powell et al., 2008; Wright et al., 2007). Interestingly, we found that their transcripts are also distributed in opposing gradients in the thalamus (Fig. S4A,B). *Ntn1* is most highly expressed at rostral-medial levels (Fig. S4A) while *Sema3a* is most highly expressed in a more caudal-lateral aspect of the thalamus and in the lateral prethalamus (Fig. S4B). In transverse *in situ* hybridization (ISH) of E13.5 controls we observed that *Ntn1* is expressed in a narrow rostral-medial thalamic population of neurons (arrow in Fig. 6A) while *Sema3a* is expressed in caudal-lateral thalamic neurons (arrows in Fig. 6B) as well as flanking the TCA bundles in the prethalamus (empty arrow in Fig. 6B).

**Figure 6.**
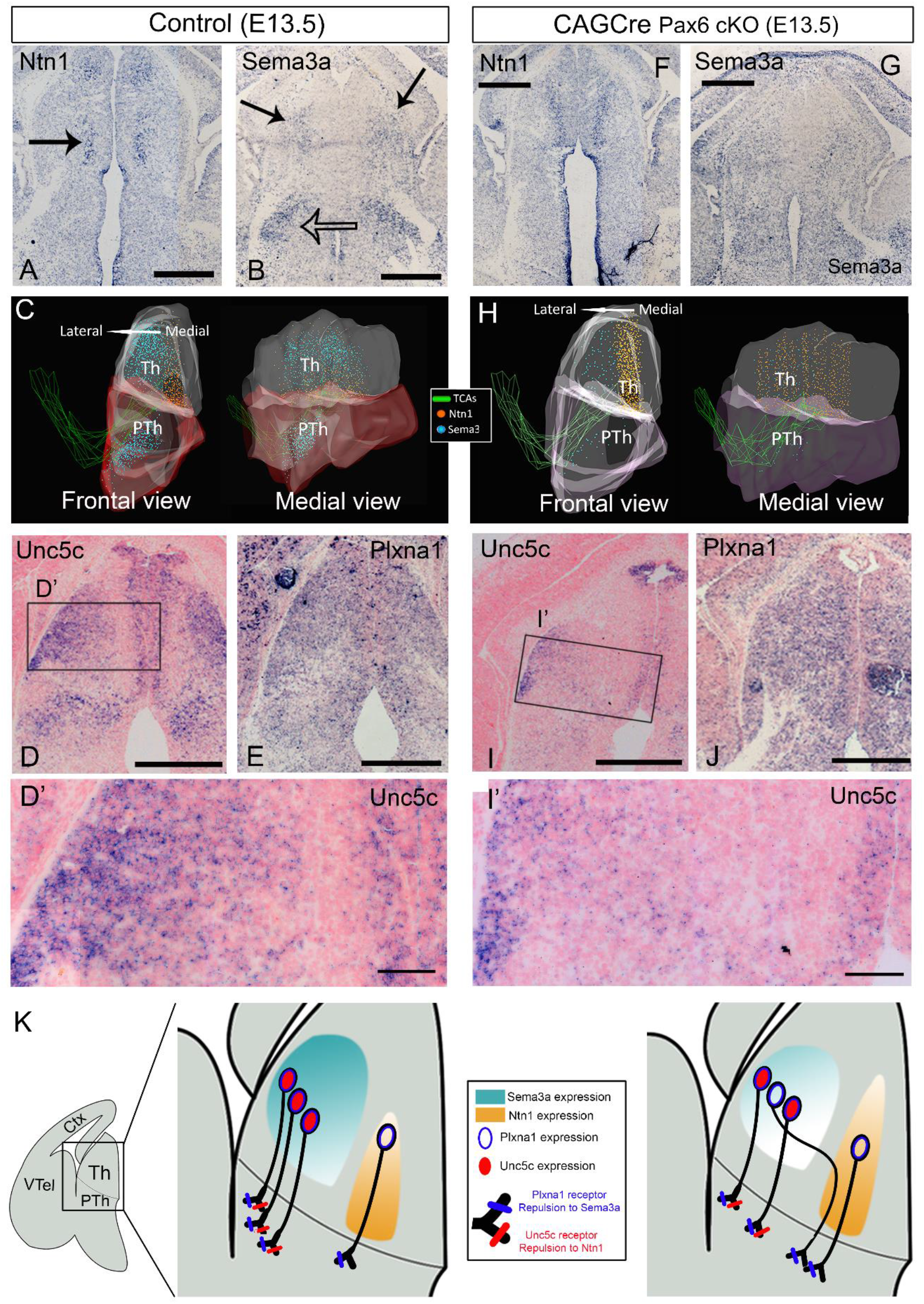
Axon guidance molecules and their receptors are altered in the diencephalon of CAG^CreER^ Pax6 cKOs. **A-I’)** Expression pattern of *Ntn1* and *Sema3a* mRNA and their receptors *Unc5c* and *Plxna1* in the E13.5 diencephalon of controls (A-E’) and CAG^CreER^ Pax6 cKOs (F-I’). C and H show two views of 3D reconstructions of the expression pattern of *Ntn1* and *Sema3a* from transverse ISH sections. **J,K)** Schematic model of the changes in *Ntn1* and *Sema3a* expression in the thalamus of controls (J) versus CAG^CreER^ Pax6 cKOs (K) and how we hypothesize might affect the guidance of different subsets of thalamic axons. In controls (J), lateral thalamic neurons express Unc5c and Plxna1, being repelled by both guidance cues, which direct their axons out from the thalamus and towards the prethalamus in a straight trajectory. In Pax6 cKOs (K) some lateral thalamic neurons lose their *Unc5c* expression but maintain their *Plxna1* expression, losing their repulsion to *Ntn1* but maintaining it for *Sema3a* and making their axons deviate towards medial thalamic regions. Scale bars: 500μm (A,B,D-E,F,G,I,J), 100μm (D’,I’). Ctx= cortex, Pth= prethalamus, Th= thalamus, vTel= ventral telencephalon. All described patterns were observed at least in three embryos belonging to three different litters for each genotype, unless otherwise stated.

To obtain a clearer three-dimensional view of these expression patterns we reconstructed them from serial, adjacent sections stained for *Sema3a, Ntn1*, Pax6 and L1 in controls (Fig. 6C, see Methods). The 3D reconstruction confirmed that *Sema3a* and *Ntn1* form opposing gradients in the normal embryonic thalamus (Fig. 6C), with *Sema3a* highest at caudal-lateral thalamic levels while *Ntn1* is highest at rostral-medial thalamic levels.

We next investigated the expression patterns of the main receptors for *Ntn1* and *Sema3a* in the thalamus of control embryos. The most interesting finding was that *Unc5c*, encoding a Ntn1 receptor mediating axonal repulsion (Leonardo et al., 1997), was expressed differentially from lateral to medial across the thalamus (Fig. 6D). Laterally, almost all cells expressed high levels of *Unc5c* whereas medially many cells did not (Fig. 6D’). *Unc5c* was largely absent from a narrow strip of cells close to and parallel with the ventricular zone. This strip coincided with the region that contained *Ntn1*-positive cells (compare Fig. 6D and A). *Plxna1*, encoding a Sema3a receptor that mediates repulsion (Rohm et al., 2000; Takahashi et al., 1999; Tamagnone et al., 1999) was found to be distributed relatively homogenously across the thalamus (Fig. 6E, S4C).

These expression patterns suggest that, whereas all thalamic axons might be repelled by Sema3a (due to their expression of *Plxna1*), only some axons might be repelled by Ntn1 (i.e. those originating laterally, which express *Unc5c*, and those *Unc5c*-expressing axons that originate medially) (Fig. 6K). This could explain why, in the grafting experiments described above, axons from lateral explants invariably navigated laterally, which would be away from medially located high levels of Ntn1. It could also explain why medial explants generated axons able to navigate on a broader front: some axons (those that express *Unc5c*) would be pushed relatively laterally by repulsion from medially expressed *Ntn1;* others (those that do not express *Unc5c*) would be able to maintain a medial trajectory through *Ntn1*-expressing territory, thereby avoiding the high levels of *Sema3a* expressed in lateral thalamus (Fig. 6K).

Other receptor-coding genes analysed (*Dcc, Unc5a, Unc5d*) showed little or no expression within the main body of the thalamus and are therefore unlikely to contribute to the navigation of thalamic axons within the thalamus (Fig. S4E,G,I).

We next asked whether the thalamic expression of *Ntn1* and *Sema3a* and their receptors change in a way that might explain the medially-directed deviation of lateral axons that we observed in the thalamus of *CAG^CreER^ Pax6* cKOs. In these embryos, we found that the medial domain of *Ntn1*-expression was retained and appeared enlarged. *Sema3a* was still expressed higher laterally, although overall levels seemed reduced (Fig. 6F,G). These patterns are reconstructed in 3D in Fig. 6H. *Ntn1* and *Sema3a* expression in the subpallium of *CAG^CreER^ Pax6* cKOs appeared to be unaffected (Fig. S4K-N). The significance of changes in the expression levels of Sema3a and Ntn1 in the thalamus of *CAG^CreER^ Pax6* cKOs was confirmed by analysing a previously published RNAseq dataset (Fig. S4C, dataset from Quintana-Urzainqui et al., 2018).

Regarding the expression of guidance receptors, fewer laterally-located neurons expressed *Unc5c* in *CAG^CreER^ Pax6* cKOs than in controls (compare Fig. 6I’ and D’). Significant numbers of *Unc5c*-negative neurons were now intermingled with *Unc5c*-positive neurons even in the most lateral thalamic tissue (Fig. 6I’). *Plxna1’s* thalamic expression pattern did not change in the absence of Pax6 (Fig. 6J, S4D), nor did that of any of the other receptor-coding genes studied (Fig. S4E-J).

As reported above, we discovered that *CAG^CreER^ Pax6* cKOs show a misrouting in a medial direction of axons from the lateral thalamus (Fig. 1), and our finding that many laterally-located thalamic neurons lose their expression of *Unc5c* in these mutants, provides a likely explanation, summarized in Fig. 6K. We propose that *Unc5c*-negative laterally-located thalamic neurons in *CAG^CreER^ Pax6* cKO thalamus would no longer be repelled from the medial thalamus by its high levels of *Ntn1*. Consequently, they would be more likely to stray, or perhaps to be pushed by relatively high lateral levels of *Sema3a*, towards a medial direction (Fig. 6K).

### Subsets of Pax6-deficient lateral thalamic axons deviate medially when confronted with control thalamus

Finally, we grafted GFP-positive lateral thalamic explants from *CAG^CreER^ Pax6* cKO embryos into the lateral thalamus of GFP-negative slices from littermate control embryos, following the same experimental paradigm as before (Fig. 7A, see Figure 5 and Methods). Our model would predict that subsets of Pax6-deficient lateral thalamic axons, presumably those that have lost their repulsion to Ntn1, would deviate towards medial areas when confronted with control thalamus. We used 11 embryos from four different litters and performed a total of 22 (bilateral) graft experiments. In all cases we observed subsets of GFP-positive axons deviating medially (Figure 7B, red arrows). To measure the navigation pattern of these axons we divided the thalamus in three equal medial-lateral sectors and quantified the percentage of GFP on each of them (Fig. 7C). For this analysis we used four explants from four different litters (5 slices per explant). We found that axons spread throughout the three areas of the thalamus (45.8% laterally, 31.8% intermedially, 22.44% medially; Fig 7C). This is in striking contrast to the almost invariably (95.7%) lateral trajectories of axons from control lateral thalamic explants grafted into control thalamic tissue (Fig. 5C-F, K). This evidence supports our model and the idea that thalamic neurons hold intrinsic information that determines the medial-lateral position of their axons when crossing the thalamus.

**Figure 7.**
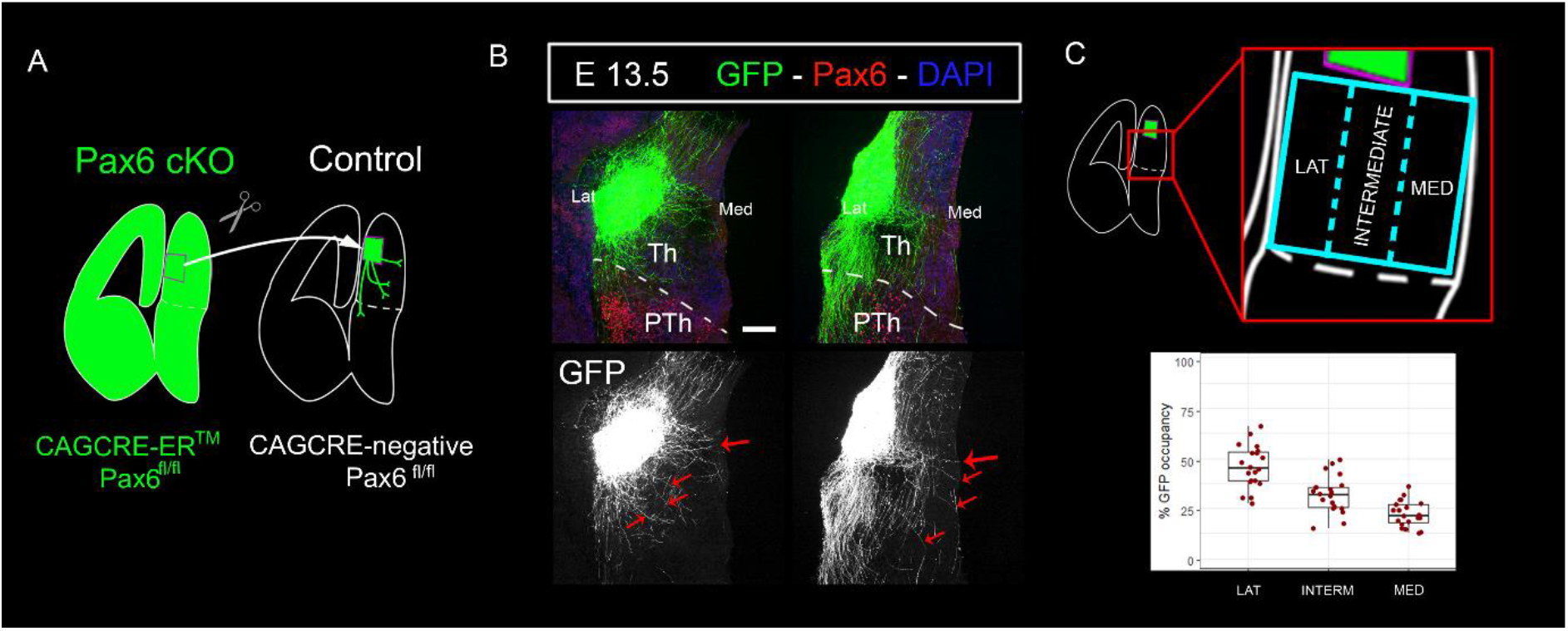
Subsets of lateral thalamocortical axons from CAG^CreER^ Pax6 cKOs deviate medially when confronted with control thalamus. **A)** Schema of the slice culture transplant experiment in which lateral thalamus from GFP-positive Pax6 cKOs embryos (CAG^CreER +^; Pax6 ^fl/fl^) was grafted into lateral thalamus of control GFP-negative littermates (CAG^CreER -^; Pax6 ^fl/fl^). **B)** Subsets of axons from Pax6 cKO lateral thalamus deviate from their normal trajectory towards medial thalamic regions (red arrows). 11 embryos from four different litters were used. A total of 22 (bilateral) graft experiments were performed. We observed medially deviated axons in all cases. **C)** Quantification (four grafts from embryos from three different litters) showed that Pax6 deficient lateral thalamic axons navigate across three delimited sectors, unlike their control counterparts which showed a strong preference for the lateral sector (Fig. 5K). Box plots show the percentage of GFP-positive axons detected on each sector of the thalamus. Five rostrocaudal sections were quantified per explant. Each dot represents an area value of a single section. Scale bar: 100μm. Lat= lateral aspect of diencephalon, Med= medial aspect of diencephalon,Th=thalamus, PTh=prethalamus

Overall, our findings indicate that mechanisms exist within the thalamus itself to ensure that its TCAs exit in an orderly manner and that these mechanisms may play an important part in the correct topographic mapping of TCAs onto the cortex.

## DISCUSSION

During embryonic development, thalamic axons undertake a long journey, navigating through several tissues before they arrive at the cortex. It is therefore crucial that their guidance is tightly regulated by mechanisms placed along the route. Previous studies have demonstrated the importance of the ventral telencephalon as an intermediate target for establishing correct topographical thalamocortical connections. Here, gradients of signalling molecules sort different subsets of TCAs towards different areas of the embryonic cortex (Antón-Bolaños et al., 2018; Bielle et al., 2011; Braisted et al., 1999; Dufour et al., 2003; Métin and Godement, 1996; Molnár et al., 2012; Vanderhaeghen and Polleux, 2004). The importance of other tissues along the route in the establishment and/or maintenance of axonal topography remained unexplored. We now show that if thalamic axons do not emerge in order from the thalamus they will connect with the wrong areas of the cortex, resulting in topographical defects. This highlights the importance of maintaining axonal order throughout the route and suggests the existence of guidance mechanisms within the diencephalon to guarantee this happens.

Our *in vitro* graft experiments demonstrated that embryonic thalamic tissue is instructive for TCAs and sorts axons according to their original medio-lateral position. We went on to find a possible guidance mechanism acting within the thalamus. The thalamus expresses *Ntn1* and *Sema3a*, some of the same guidance molecules known to guide TCAs in the ventral telencephalon (Bielle et al., 2011; Powell et al., 2008; Wright et al., 2007). What is more, there is an interesting correspondence between the regions expressing each of those molecules in the thalamus and in the ventral telencephalon. TCAs that emerge and navigate through the *Sema3a*-high region of the thalamus (lateral-caudal thalamus, dLGN) are steered towards the *Sema3a*-high region in the ventral telencephalon, while axons that emerge and navigate through the *Ntn1*-high region of the thalamus (ventral-medial thalamus,VMP) are sorted towards *Ntn1*-high regions in the ventral telencephalon. This suggests that each subset of thalamic axons maintains the expression of the same combination of axon guidance receptors along the route and therefore show the same chemotactic response when confronting gradients of signalling cues. Likewise, it indicates that the same gradients of guidance molecules are re-used at different levels of the thalamocortical pathway to maintain topographic order.

The chemotactic behaviour of TCAs with respect to *Sema3a* and *Ntn1* gradients in the thalamus and ventral telencephalon can be explained by our observations of the expression of Sema3a and Ntn1 receptors in developing thalamus. We show that all thalamic neurons seem to express homogeneous levels of *Plxna1*, a receptor mediating repulsion to Sema3a (Rohm et al., 2000; Takahashi et al., 1999; Tamagnone et al., 1999), while Unc5c, a receptor mediating repulsion to Ntn1 (Leonardo et al., 1997), was found to be expressed in a lateral-high medial-low gradient. According to these observations, we propose a model in which complementary expression patterns of Sema3a and Ntn1 can establish topographical order on TCAs by a mechanism of double repulsion, in which all thalamic axons have the potential to be repelled by Sema3a but only lateral axons are additionally repelled by Ntn1. It is possible that laterally-derived axons experience stronger Ntn1 repulsion the more lateral they are. Thus, lateral thalamic axons prefer to navigate through Sema3a-high, Ntn1-low regions because they might be more strongly repelled by Ntn1 than by Sema3a. Axons located in intermediate regions of the thalamus express lower levels of Unc5c; thus they might be equally repelled by Sema3a and Ntn1 and chose to navigate across regions with moderate levels of both signalling cues. Likewise, medial axons are only repelled by Sema3a and neutral to Ntn1: therefore they chose to navigate through Sema3a-low, Ntn1-high areas.

Supporting this model are the experiments showing that TCAs are repelled by Nnt1 (Bielle et al., 2011; Bonnin et al., 2007; Powell et al., 2008) and thalamic growth cones show retraction in the presence of Sema3a (Bagnard et al., 2001). Moreover, Wright and colleagues reported that in mice harbouring a mutation that makes the axons non-responsive to Sema3a, axons from the ventrobasal (VB) thalamic nucleus were caudally shifted and target the visual cortex instead of the somatosensory cortex (Wright et al., 2007). Our double repulsion model satisfactorily explains this phenotype. The VB nucleus is located in an intermediate thalamic region that would contain substantial number of Unc5c-positive neurons. In those mutants, VB axons lose their repulsion to Sema3a but many would still be repelled by Ntn1, and therefore would steer towards Sema3a-high, Ntn1-low regions both in the thalamus and the ventral telencephalon.

The behaviour of thalamic axons in our explant experiments using donor tissue from controls and CAG^CreER^ Pax6 cKOs, also support the model.

Other molecules known to form gradients and guide TCAs in the ventral telencephalon, like Slit1 or Ephrin A5 (Bielle et al., 2011; Dufour et al., 2003; Molnár et al., 2012; Seibt et al., 2003; Vanderhaeghen and Polleux, 2004) were not analysed in this study. It remains to be tested whether these molecules and their receptors are expressed in the thalamus in a gradient fashion and if they follow the same rules proposed in our model.

It is important to highlight that in this study we only considered the medio-lateral axis of the main thalamic body, but the same or other guidance cues and receptors probably function in other directions. For example, work in mice showed that Unc5c (Bonnin et al., 2007) and DCC (Powel et al., 2008) are also highly expressed in the rostral thalamus, at a level we did not cover in our expression and tracer analyses.

Other important question is how Pax6 inactivation leads to deficits in thalamic organization, since Pax6 is only expressed in progenitors. We show here and in a previous paper that thalamic patterning does not seem to be largely affected in CAG^CreER^ Pax6 cKOs (Fig. S1 and Quintana-Urzainqui et al., 2018) and that the main changes in gene expression in postmitotic neurons are related with axon guidance molecules. One possibility is that Pax6 expression in thalamic progenitors affects transcriptional programmes that indirectly translates into actions on the postmitotic expression of certain axon guidance molecules.

Finally, our results have given interesting new insights into the development and the importance of the pioneer axons from the prethalamus to the thalamus. First, we found that the prethalamic neurons extending pioneer axons to the thalamus belong to a particular lineage, the Gsx2-lineage and not the Zic4-lineage. Second, although disturbing the prethalamus-to-thalamus pioneers did not stop TCAs reaching the cortex without any topographic error, it did cause them to fasciculate prematurely as they crossed the prethalamus. Growing axons often increase their fasciculation when they cross regions that are hostile to their growth. Our and other studies have shown that the prethalamus also expresses guidance cues with potential to exert a repulsive response of TCAs (Ono et al., 2014), thus it is possible that interactions between the developing TCAs and prethalamic pioneers somehow helps the passage of the TCAs across this region. Further work is needed to discover what the consequences are if this help is unavailable.

## Material and Methods

### Mice

All animals (*Mus musculus*) were bred according to the guidelines of the UK Animals (Scientific Procedures) Act 1986 and all procedures were approved by Edinburgh University’s Animal Ethics Committee.

The day the vaginal plug was detected was considered E0.5

For conditional inactivation of Pax6, we used a tamoxifen-inducible Pax6^loxP^ allele (Simpson et al., 2009) and a RCE:LoxP EGFP Cre reporter allele (Sousa et al., 2009) and we combined them with different Cre lines. To generate a deletion of Pax6 throughout the embryo, we used lines carrying a CAGGCre-ER^™^ allele (Hayashi and McMahon, 2002; Quintana-Urzainqui et al., 2018). As controls, we used both wild type and *CAG^CreER^ Pax6^fl/+^* littermate embryos since the latter express normal levels of Pax6 protein, almost certainly because of a feedback loop that compensates for a deletion in one allele by increasing the activity of the other (Caballero et al., 2014; Manuel et al., 2015).

To inactivate Pax6 in different parts of the prethalamus we used either a Gsx2-Cre (Kessaris et al., 2006) or the Zic4-Cre allele (Rubin et al., 2011). For cortex-specific deletion of Pax6, we used Emx1Cre-ER^T2^ (Kessaris et al., 2006).

The DTy54 YAC reporter allele (Tyas et al., 2006) was combined with the Gsx2-Cre allele to generate Gsx2Cre; Pax6^loxP/loxP^ embryos expressing tauGFP in cells in which the Pax6 gene is active.

Embryos heterozygous for the Pax6^loxP^ allele (Pax6^fl/+^) were used as controls since previous studies have shown no detectable defects in the forebrain of Pax6^fl/+^ embryos (Simpson et al., 2009). Embryos carrying two copies of the floxed Pax6 allele (Pax6^fl/fl^) were the experimental conditional knock-out (cKO) groups.

For thalamic explant experiments, we generated litters containing GFP-positive and negative embryos by crossing a line of heterozygous studs for a constitutively active form of CAGGCre-ER^™^ allele and the and a RCE:LoxP EGFP Cre reporter allele with wild type females. In the second set of explant experiments, we performed crossings to generate litters containing GFP-positive Pax6 cKO embryos (CAGCRE^ER TM^; Pax6^fl/fl^), and GFP-negative control embryos (homozygous for the Pax6 floxed allele but not expressing the CRE allele). Both embryos carried a RCE:LoxP EGFP allele reporting Cre activity and allowing us to select the embryos within the same litter and visualizing Pax6 cKO axons in the explants

The day the vaginal plug was detected was considered E0.5. Pregnant mice were given 10mg of tamoxifen (Sigma) by oral gavage on embryonic day 9.5 (E9.5) and embryos were collected on E12.5, E13.5, E14.5, E15.5, E16.5 or E18.5 For the DiI and DiA tracing experiments, wild type embryos (CD1 background) were additionally used as controls.

### Immunohistochemistry

Embryos were decapitated and fixed in 4% paraformaldehyde (PFA) in phosphate buffered saline (PBS) overnight at 4°C. After washes in PBS, heads were cryoprotected by immersion in 30% sucrose in PBS, embedded in OCT Compound and sectioned using a cryostat at 10μm.

Cryo-sections were let to stabilize at room temperature for at least 2 hours and then washed three times in PBST (1X PBS with 0.1% Triton X-100, Sigma). To block endogenous peroxidase, sections were treated with 3% H_2_O_2_ for 10 minutes. After PBS washes, antigen retrieval was performed by immersing the sections in Sodium Citrate buffer (10mM, pH6) heated at approximate 90°C using a microwave for 20 minutes. Sections were then incubated with the rabbit polyclonal anti-Pax6 (1:200, BioLegend Cat # 901302) overnight at 4°C. The secondary antibody (goat anti-rabbit bioninylated, 1:200, Vector laboratories Cat # BA-1000) was incubated for 1 hour at room temperature followed by a 30-minute incubation with Avidin-Biotin complex (ABC kit, Vector laboratories Cat # PK6100). Finally, diaminobenzidene (DAB, Vector Laboratories, Cat # SK4100) reaction was used to obtain a brown precipitate and sections were mounted in DPX media (Sigma-Aldrich, Cat # 06522).

For immunofluorescence, cryosections were incubated overnight at 4°C with the following primary antibodies: rat monoclonal anti-Neural Cell Adhesion Molecule L1 (1:500 Millipore, Cat # MAB5272, clone 324, RRID:AB_2133200), rabbit polyclonal anti-Pax6 (1:200, BioLegend Cat # 901302; RRID:AB_2565003), goat polyclonal anti-GFP (1:200, Abcam Cat # ab6673, RRID:AB_305643), rabbit polyclonal anti-GFP (1:200, Abcam Cat # ab290, RRID:AB_303395). The following secondary antibodies from Thermo Fisher Scientific were incubated at room temperature for one hour: Donkey anti-rat Alexa^488^ (1:100, Thermo Fisher, Cat # A-21208), Donkey anti-rat Alexa^594^ (1:100, Thermo Fisher, Cat # A-21209), Donkey anti-rabbit Alexa^568^ (1:100, Thermo Fisher, A10042), Donkey anti-rabbit Alexa^488^ (1:100, Thermo Fisher Cat # R37118), Donkey anti-goat Alexa^488^ (1:100, Invitrogen, Cat # A11055). Sections were counterstained with DAPI (Thermo Fisher Scientific, Cat # D1306) and mounted in ProLong Gold Antifade Mountant (Thermo Fisher Scientific, Cat # P36930).

### *In situ* hybridization

*In vitro* transcription of digoxigenin-labelled probes was done with DIG RNA-labeling kit (Sigma-Aldrich, Cat # 11175025910). The following digoxigenin-labelled probes were synthetized in the lab from cDNA: Ntn1 (kindly donated by Dr Thomas Theil; forward primer: CTTCCTCACCGACCTCAATAAC, reverse primer: GCGATTTAGGTGACACTATAGTTGTGCC TACAGTCACACACC), Sema3a (forward primer: ACTGCTCTGACTTGGAGGAC, reverse primer: ACAAACACGAGTGCTGGTAG), Plxna1 (forward primer: GACGAGATTCTGGTGGCTCT, reverse primer: CATGGCAGGGAGAGGAAGG), DCC (forward primer: AACAGAAGGTCAAGCACGTG, reverse primer: CAATCACCACGACCAACACA), Unc5a (forward primer: CTGTCAGACCCTGCTGAGT, reverse primer: GGGCTAGAGTTCGCCAGTC), Unc5d (forward primer: GGACAGAGCTGAGGACAACT, reverse primer: GTATCAAACGTGGCGCAGAT). Unc5c probe was kindly donated by Dr. Vassiliki Fotaki, University of Edinburgh, UK and Dr Suran Ackerman, UCSanDiego, USA). Cryosections were processed for *in situ* hybridization (ISH) using standard protocols. Some slides were counterstained for nuclear fast red (Vector Laboratories, Cat# LS-J1044-500).

### Axon Tract Tracing

For cortical injections, brains were dissected between E15.5 and E18.5 and fixed in 4% PFA in PBS at 4°C for at least 48 hours. After washes in PBS, filter paper impregnated in DiI (NeuroVue Red, Molecular Targeting Technologies, Cat # FS-1002) and DiA (NeuroVue Jade, molecular Targeting Technologies, Cat # FS-1006) was inserted approximately in the somatosensory and visual areas of the cortex, respectively. Brains were incubated at 37°C in PBS for 4 weeks to allow the diffusion of the tracers.

For thalamic injections in fixed tissue, embryos were dissected at E13.5 and fixed overnight in 4%PFA in PBS at 4°C. After PBS washes, brains were cut in half at the midline and DiI was inserted in the thalamus using a fine probe. Brains were incubated for 1 week in PBS at 37°C.

Brains were then cryoprotected in 30% sucrose, embedded in OCT Compound and sectioned in a cryostat at 30μm. Sections were counterstained with DAPI diluted 1:1000 in distilled water.

For thalamic injections in non-fixed tissue, we applied neurobiotin (Vector Laboratories, Cat # SP-1120), and amino derivative of biotin used as an intracellular label for neurons. The tracer in powder was held at the tip of an entomological needle (00) and recrystallized using vapour from distilled water. Brains were cut in half and the crystal was inserted in the thalamus. Brains were then immersed in continuously oxygenated Ringer (124mM NaCl, 5mM KCl, 1.2mM KH2P04, 1.3mM MgSO4 7H2O, 26mM NaHCO3, 2.4mM CaCl2 2H2O, 10mM glucose) and incubated overnight at RT. The tissue was fixed in 4% PFA in PBS overnight at 4°C, washed in PBS, cryoprotected in 30% sucrose and sectioned in a cryostat at 10μm. Neurobiotin was visualized by incubating the sections with either Strep^488^ or Strep^546^.

### Thalamic explants and slice culture

E13.5 embryos were dissected, embedded in 4% low melting temperature agarose (Lonza, Cat # 50100) and sectioned in a vibratome to produce 300μm-thick coronal slices. Lateral or medial thalamic explants were dissected from slices belonging to GFP-positive embryos and transplanted into equivalent rostral/caudal slices belonging to GFP-negative embryos (see schemas in Fig. 5). The thalamus and its different medio-lateral regions were recognized under the dissecting scope by anatomical landmarks. Slices were then cultured for 72 hours in floating membranes (Whatman nuclepore track-etched membranes, Cat# WHA110414) over serum-free Neurobasal medium (Thermo Fisher Scientific, Cat# 21103049) in 60mm center well organ culture dishes (Falcon, Cat# 353037). Cultures were fixed in 4% PFA overnight at 4°C, cryoprotected in 30% sucrose and cryosectioned at 10μm to be processed for immunofluorescence. All grafts were positioned in direct contact with presectioned host tissue to avoid pial growth.

We performed a total of 31 transplant experiments. In the first set of experiments, we grafted control thalamus into control tissue. We used embryos from four different litters to a total of nine different transplants (four using lateral thalamus and five using medial thalamus as donor tissue). In the second set of experiments, we grafted Pax6 cKO lateral thalamus into control thalamus, using 11 embryos from four different litters and a total of 22 individual (bilateral) transplants.

### Image analysis and quantification of thalamic area occupied by DiI and DiA

Images of transverse sections were analysed blind using Fiji Software (Schindelin et al., 2012) for seven E15.5 embryos belonging to three different litters (four controls and three *CAG^CreER^ Pax6* cKOs). For each embryo, we analysed at least 5 sections at different rostral-caudal levels. For each section, we isolated the DAPI channel and defined the dLGN and VP nucleus as regions of interest (ROIs) blind to the other channels (DiI and DiA). We next measured the area occupied by DiI and DiA within each ROI, defining positive label using the automated thresholding function in Fiji software set for “MaxEntropy”. Data from all sections and genotypes was statistically assessed to test the effects of Pax6 inactivation on the area occupied by each dye on each nucleus. Data was fitted to a mixed linear model using the lmer() function from lme4 R package (Bates et al., 2015). “DiI in dLG”, “DiI in VP”, “DiA in dLG” and “DiA in VP” were set as dependant variables, with “genotype” as a fixed effect and “litter:embryo” factors as nested random effects. P-values of fixed effects were obtained using the Anova() function from car package (Fox and Weisberg, 2011).

### Quantification of thalamic explant experiments

Images of transverse sections of our explants were analysed using Fiji Software (Schindelin et al., 2012). We quantified a total of ten culture thalamic explants (three from lateral control donor tissue, three from medial control donnor tissue and four from lateral Pax6 cKO tissue) from seven different litters, using five slices per explant corresponding to different rostral-caudal levels of the explants. For each section we first divided the thalamus into three equal medial-lateral sectors (defined as ROIs, see Figure 5), avoiding the area occupied by the explant itself. We then isolated the GFP channel and measured the area occupied by GFP-positive elements within each ROI, defining positive label using the automated thresholding function in Fiji software set for “MaxEntropy”. To test for the differences of GFP occupancy in the three different areas (lateral, intermedial, medial), the data was fitted to a mixed linear model using the lmer() function from lme4 R package (Bates et al., 2015).

For the DiI and thalamic explant quantifications, data was presented as box plots representing the median value and the distribution of the quartiles. Individual datapoints representing measurements of each slice were plotted overlying the box plots.

### Quantifications of numbers of axons and bundle width

Images were blinded analysed for at least three E13.5 embryos for each condition belonging to three different litters. We positioned three lines across the prethalamus: (1) at the thalamic-prethalamic border (Th-PTh), guided by prethalamic expression of Pax6; (2) at a lower prethalamic position (low-PTh), guided by the end of Pax6 prethalamic expression; and (3) at the midpoint position between the two other lines (mid-PTh). We generated a L1 intensity profile using Fiji Software (Schindelin et al., 2012). Intensity profiles were then processed by tracing a line at an arbitrary (but constant for all quantifications) intensity level and quantifying the number and width of bundles crossing the line. Statistical significance was assessed applying two-tailed unpaired Student’s t-test and N=3.

### 3D reconstruction

We used Free-D software (Andrey and Maurin, 2005) to reconstruct the structure of thalamus and prethalamus from transverse slices stained with DAPI and antibodies against L1 and Pax6 to reveal the thalamocortical tract and the limits of the diencephalic structures, respectively. The thalamus territory was recognisable by an intense DAPI staining and Pax6-negative mantle zone, contrasting with prethalamus and pretectum, which express high levels of Pax6 in the postmitotic neurons. The location of the signalling molecules was included in the model by comparison of transversal and sagittal sections stained for Ntn1 and Sema3a and their adjacent sections processed for Pax6 and L1 with the sections used to build the model scaffold. Dots are representation of staining density. We used sagittal and transverse sections from four embryos from three different litters.

### Microscopy and imaging

ISH and IHQ images were taken with a Leica DMNB microscope coupled to a Leica DFC480 camera. Fluorescence images were taken using a Leica DM5500B automated epifluorescence microscope connected to a DFC360FX camera. Image panels were created with Adobe Photoshop CS6.

## Acknowledgements

We thank Dr Martine Manuel for her invaluable help with the generation of mice lines for the explant experiments and Dr Vassiliki Fotaki for providing the Unc5c probe.

